# The Gut-Brain Axis Shapes Cognitive-Emotional Processing: Evidence for Attentional Avoidance of Bloating Cues in IBS

**DOI:** 10.1101/2025.09.01.673534

**Authors:** Reyhaneh Akbari, Fateme Dehghani-Arani, Mohsen Honar, Nazila Shahmansouri, Ehsan Rezayat

**Affiliations:** Department of Psychology, Faculty of Psychology and Education, University of Tehran, Tehran, Iran; Department of Cognitive Sciences, Faculty of Psychology and Education, University of Tehran, Tehran, Iran; Tehran Heart Center, Tehran University of Medical Sciences, Tehran, Iran

**Keywords:** Irritable bowel syndrome, attentional bias, dot-probe, somatic threat, gut-brain axis

## Abstract

**Background:** Irritable bowel syndrome (IBS) is a functional gastrointestinal disorder characterized by significant gut-brain interactions and various symptoms such as abdominal pain, bloating, and altered bowel habits. Cognitive deficits in IBS have been linked to attentional biases, particularly towards somatic and symptom-related cues. Despite increasing interest in understanding these cognitive processes, the specific patterns of attentional biases and their relationship with anxiety in IBS remain inadequately explored.

**Method:** This study employed a dot-probe task to compare attentional biases toward somatic (bloating, pain) and social threat cues (angry/disgusted faces) in 15 patients with IBS, 15 individuals with high anxiety (HA), and 15 healthy controls (HC). Participants completed two tasks (Body/Face) with 500 ms stimulus exposure. Attentional bias indices (RT incongruent – RT congruent) were analyzed using mixed-design ANOVAs, controlling for anxiety (STAI) and IBS symptom severity (IBS-SSS).

**Results:** Patients with IBS exhibited significant avoidance of bloating-related stimuli compared to HC (p =.013), but not pain-related cues. Anxiety modulated attentional processing in IBS (p =.015) and HC (p =.013). HA showed no significant biases. IBS patients demonstrated slower overall reaction times than HC (p=.024), suggesting cognitive load. A positive IBS severity-anxiety correlation was observed in IBS (r = 0.574, p =.020).

**Conclusions:** The findings demonstrate that IBS is associated with selective avoidance of bloating-related stimuli, driven by a maladaptive interplay between hyper-precise symptom expectations and interoceptive noise—a mechanism distinct from anxiety-related attentional patterns. While anxiety amplifies this avoidance, it does not independently account for the cognitive profile of IBS. These results underscore the gut-brain axis’ role in shaping cognitive-emotional processing.

## 1. Introduction

Irritable bowel syndrome (IBS) is a disorder of gut–brain interactions (Sperber, 2021) characterized by altered bowel habits and a cluster of symptoms including abdominal pain, constipation, feeling of urgency, and incomplete evacuation for at least 3 months (Drossman & Hasler, 2016). The estimated global prevalence of this condition is between one in 11 and one in 26 people globally, depending on the definition used (Oka et al., 2020). Although traditionally it has been considered that there is no clear disease or inflammatory marker for IBS (Holtmann et al., 2016), factors including microbiota, low-grade inflammation, and alterations to gut functioning and sensitivity play a role in its pathophysiology (Canakis et al., 2020).

Despite occurring in all age groups, the prevalence of IBS in people over 50 years of age is 25% lower than in people aged 50 years or younger (Lovell & Ford, 2012). IBS can impair quality of life, sometimes leading to unemployment, limitations in activity and social functioning (Canavan et al., 2014; Mönnikes, 2011). Furthermore, anxiety are common in patients with IBS, affecting one-third of patients witth IBS (Mönnikes, 2011). Interestingly, in about half of individuals with IBS, the condition seems to develop first, with anxiety emerging later. This suggests that disruptions in gut function may be the primary drivers of anxiety (Jones et al., 2012; Koloski et al., 2016). These findings have led to the conceptualization of IBS as a primary disorder of the gut-brain axis, where the brain is believed to influence gut-related symptoms (Quigley, 2018). Gut-brain axis is a constant two-way communication between the brain and the digestive system, involving neural, endocrine, and metabolic signals. Any disruption in the digestive system (referred to as the “little brain” due to its effects (Fichna & Storr, 2012; Quigley, 2018) can negatively impact the brain and lead to cognitive impairment (Kennedy et al., 2012; Weaver et al., 2016). Emerging evidence suggests that individuals with IBS may exhibit cognitive impairments, particularly in attentional bias (Lam et al., 2019; Wong et al., 2019). Attentional bias—the tendency to prioritize the processing of certain types of stimuli over others (Azriel & Bar-Haim, 2020)—has been observed in various conditions associated with heightened sensitivity to stress and anxiety (Cisler & Koster, 2010; Van Bockstaele et al., 2014). In IBS, this bias may manifest as a hypervigilance to negative and symptom-related stimuli. People with IBS demonstrate a confirmatory bias towards emotionally negative and pain-related information (Chapman & Martin, 2011; Gomborone et al., 1993; Phillips et al., 2014)

Early studies using Stroop and memory tasks revealed that patients with IBS process pain-related and emotionally negative information more automatically, showing faster engagement and more false-positive errors compared to healthy controls (Afzal et al., 2006; Chapman & Martin, 2011; Gomborone et al., 1993). Subsequent neurophysiological studies provided biological evidence for these cognitive biases (Ejova et al., 2021; Hong et al., 2016). Key psychological traits— particularly neuroticism, anxiety and symptom severity—have been shown to correlate with and potentially amplify these attentional patterns (Hubbard et al., 2015; Phillips et al., 2014). Recent work suggests these biases may represent stable cognitive markers of IBS rather than temporary states (Akbari et al., 2025; Wong et al., 2019).

While previous research has primarily focused on pain-related attentional biases in IBS using verbal stimuli (Afzal et al., 2006; Chapman & Martin, 2011; Posserud et al., 2009; Tkalcic et al., 2014), less is known about how patients process bloating-related cues, despite bloating being one of the most distressing symptoms (Chiarioni, 2019; Kanazawa et al., 2016; Schmulson & Drossman, 2017). Additionally, facial expressions of anger and disgust are particularly relevant for several reasons. First, anger represents a social threat cue that may trigger stress responses in patients with IBS, who often report heightened social anxiety (Zamani et al., 2019). Negative social interactions are more strongly associated with IBS outcomes than supportive relationships - conflict and adverse exchanges in social relationships correlate with increased stress and impaired quality of life in patients with IBS (Lackner et al., 2013). Second, disgust - linked to visceral sensitivity - may resonate more strongly with patients with IBS due to their heightened interoceptive awareness (Longarzo et al., 2017) and the condition’s frequent association with disgust sensitivity (Formica et al., 2022) likely related to altered processing of sensory stimuli along the brain-gut axis (Carpinelli et al., 2023). Comparing responses to emotional (facial) versus somatic (bodily) stimuli can clarify whether attentional biases in IBS are disorder-specific (i.e., tied to gastrointestinal symptoms) or reflect a broader tendency toward negative emotional processing that includes social threat perception. This gap in the literature is particularly notable given the ongoing debate about the most appropriate stimulus modality for assessing attentional biases. There is evidence that using images over words as stimuli in the dot-probe task because their presumed higher ecological validity compared to words (Staugaard, 2009). As highlighted in a recent meta-analysis by Abudoush et al. (2023), the apparent advantage of verbal stimuli may simply reflect their predominant use in attentional bias research, with relatively fewer studies employing pictorial stimuli despite their potential ecological validity.

The dot-probe task has been widely used in cognitive psychology to assess attentional biases toward threat-related stimuli (MacLeod et al., 1986). In this paradigm, participants are presented with paired stimuli (e.g., one threatening and one neutral), followed by a probe that appears in the location of one of them. Faster reaction times to probes replacing threat-related stimuli indicate attentional vigilance, while slower responses suggest avoidance. Currently, the dot probe task is used widely to measure attentional bias toward threat (Cannito et al., 2020; Wang et al., 2021; Wieser & Keil, 2020) Given that patients with IBS exhibit heightened sensitivity to symptom-related and negative information (Larsson et al., 2012; Phillips et al., 2014), the dot-probe task offers a well-validated method to investigate whether these biases extend to both facial expressions (angry, disgusted) and somatic stimuli (bloating, pain) (Price et al., 2015).

Despite advancements in the field, it remains unclear how different types of stimuli (e.g., facial expressions vs. bodily states) may influence attentional biases in patients with IBS compared to individuals with high anxiety (HA) and healthy controls (HC). The lack of control for factors such as anxiety and IBS symptoms severity also make it difficult to determine their specific contributions to the development and persistence of IBS symptoms. A key unresolved question is how much of the attentional bias observed in patients with IBS stems from dysfunction of the brain-gut axis—and is thus directly related to IBS itself—versus how much is influenced by comorbid factors such as anxiety. This gap in understanding highlights the need for more nuanced research to disentangle the complex interplay between cognitive, emotional, and physiological mechanisms in IBS.

Thus, the relationship between brain-gut interactions and attentional biases in IBS remains poorly understood. To address these gaps, our study employs a dot probe task divided into two parts: one presenting images of angry and disgusted faces, and the other showing bodies in states of bloating or pain — patients with IBS, individuals with high anxiety (non-clinical), and healthy controls— to better understand the mechanisms underlying attentional biases in IBS and their potential links to anxiety and brain-gut dysfunction.

Our hypothesis posits that the disorder of brain-gut interactions in patients with IBS will result in a unique pattern of attentional bias, distinct from that observed in individuals with high anxiety and healthy controls. Specifically, we predict that patients with IBS will show a heightened attentional bias toward bodily states of bloating and pain. In contrast, individuals with high anxiety may exhibit a broader attentional bias toward both facial and bodily threat cues, while healthy controls are expected to show minimal bias across all conditions.

## 2. Methods

### 2.1. Participants

A total of 17 individuals diagnosed with irritable bowel syndrome (IBS), 15 individuals with high anxiety (HA) and 15 healthy controls (HC) were included in the current study. Patients with IBS were recruited via online advertisements distributed through social media platforms, IBS-specific forums, and university-affiliated research portals. Participants completed an initial screening survey by a physician followed by a structured clinical interview to validate diagnosis and exclude comorbid gastrointestinal or psychiatric conditions. Groups were matched for sex, age, and education level to ensure demographic comparability. Of the 17 patients with IBS initially recruited, two were excluded due to comorbidities (endometriosis and prior colonic surgery). HC and participants with HA participants were recruited using parallel methods. As a result, 15 patients with IBS (age, 30.19 ± 7.3 years; 10 females and 5 males; education level 18.15± 1.3 years), 15 individual with HA (age, 28.31 ± 7.2 years; 10 females and 5 males; education level 17± 1.63 years), and 15 HCs (age, 29.07 ± 5.5 years; 10 females and 5 males; education level 18.13 ± 1.76 years) were further assessed. All participants were right-handed, with normal or corrected-to-normal visual. Inclusion criteria were: (a) primary education level or above; (b) aged between 18 and 50 years; For the IBS group specifically, an additional inclusion criterion was: (c) a confirmed diagnosis of IBS by a specialist physician and fulfillment of the ROME IV criteria. Individuals were excluded from the study if they had (a) a history of severe psychiatric disorders (e.g., schizophrenia, bipolar disorder, or major depressive disorder); (b) a history of organic gastrointestinal diseases (e.g., inflammatory bowel disease, celiac disease, or colorectal cancer); (c) a history of abdominal surgery (except for appendectomy or cholecystectomy performed more than one year prior); (d) abuse of alcohol or psychoactive substances in the past year; (e) current pregnancy or breastfeeding in women; (f) use of medications that significantly affect gastrointestinal motility or sensitivity (e.g., opioids, anticholinergics) within the past four weeks; or (g) participation in another clinical trial or intervention study within the past three months.

Individuals who are placed in the group with high anxiety in this study are those who, through an interview, acknowledge experiencing high levels of anxiety in their lives and were subsequently assessed using the Spielberger State-Trait Anxiety Inventory (STAI) questionnaire. Based on the cutoff score of this questionnaire, individuals with an anxiety score higher than 95 were classified as having high anxiety and placed in the anxiety group (Vigneau & Cormier, 2008). It is important to note that individuals in the group with high anxiety do not have any clinical anxiety disorders (such as generalized anxiety disorder, social anxiety disorder, etc.). Individuals with high anxiety and HC had no history of IBS, brain illness history, psychiatric disorders or medications, as confirmed by a structured diagnostic interview.

A priori power analysis (GPower 3.1.9.2) for the critical Group × Stimulus Type × Dot Type interaction in a mixed ANOVA indicated that N=27 total (9 per group). Based on prior attentional bias studies in IBS (Lam et al., 2019), we targeted 84% power to detect medium effects (f=0.25, α=0.05). Our final sample (N=15 per group, 45 total) exceeds this requirement, providing >95% power for similar effects.

### 2.2. Measures

The Persian version of Spielberger State-Trait Anxiety Inventory (STAI) questionnaire was administered to assess anxiety levels in individuals with high anxiety (Abdoli et al., 2020). The STAI is a widely used self-report measure that evaluates both state anxiety (temporary feelings of anxiety) and trait anxiety (general predisposition to anxiety). Participants rated their agreement with statements on a 4-point Likert scale, with higher scores indicating greater anxiety. This measure was used to confirm the high anxiety status of participants in the group with high anxiety and to ensure that healthy controls (HCs) and did not exhibit elevated anxiety levels (Spielberger et al., 1971).

Additionally, the IBS-Symptom Severity Scale (IBS-SSS), developed in 1997 (Francis et al., 1997; Spiegel et al., 2009), was used to assess symptom severity in patients with IBS. This widely used measure includes Visual Analog Scales (VAS), open-ended and binary questions, and a diagram of the abdomen. It comprises nine items with several sub-items, four of which focus on abdominal pain, including pain intensity, frequency, location, and its relation to stool patterns. The IBS-SSS is a validated and frequently employed tool in IBS trials and cohort studies, providing a comprehensive assessment of symptom severity (Williams et al., 2014).

#### 2.2.1. Dot Probe Task

This study incorporated two computerized dot probe tasks (Face and Body) to assess attentional biases (figure 1). In the dot probe paradigm, —one threat-related and one neutral—are presented simultaneously at different spatial locations on a computer screen (McLeod, Matthews, & Tata, 1986). After a brief interval, the cues disappear, and a target probe (e.g., a dot) appears at one of the two previous locations.

**Figure 1.**
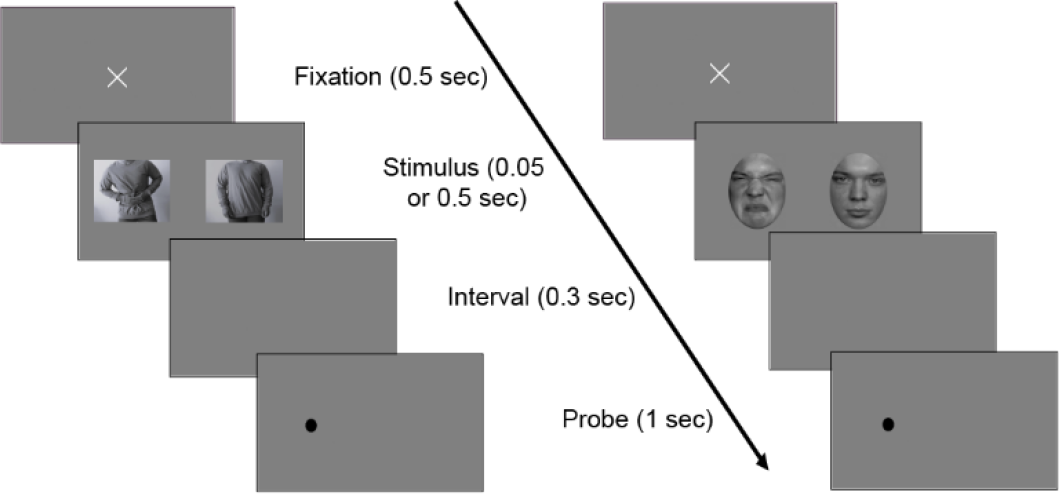
Trial structure and experimental conditions in the visual dot probe task. The task begins with a fixation period of 500 ms, followed by the simultaneous presentation of a threat-related and a neutral stimulus for 500 or 50 ms. After a 300 ms interval, a probe (e.g., a dot) appears at one of the previous stimulus locations for 1000 ms. Participants are required to respond to the probe’s location as quickly and accurately as possible. In the Body Task (left sequence), the stimuli consist of images of bodies, while in the Face Task (right sequence), the stimuli are facial expressions. Congruent trials occur when the probe replaces the threat-related stimulus, while incongruent trials occur when the probe replaces the neutral stimulus. Reaction times on these trials are used to assess attentional bias toward or away from threat-related information.

##### Face Task

Eighteen black-and-white facial images (6 disgust, 6 anger, 6 neutral) were selected from the validated Dynamic FACES database, featuring six individuals (3 males, 3 females) (Holland et al., 2019). The stimuli represented a balanced age distribution, including both young adults and middle-aged individuals. Non-facial contours and hair were removed using Adobe Photoshop, and images were resized to fit an oval-shaped frame. The gender ratio of each emotion type was 1:1.

##### Body Task

Eighteen body images (6 neutral, 6 lower abdominal pain, 6 bloating) were captured by the researcher using anonymized photographs of volunteers. The sample included individuals of varying ages (young adults to middle-aged) and diverse body types (BMI range: 18.5–30 kg/m^2^) to ensure representativeness. Volunteers provided explicit written consent for image use, with all identifiable features (e.g., faces) removed to ensure anonymity. Photographs were taken using a fixed camera distance (1.5m), uniform lighting, and neutral backgrounds to standardize posture and expression. The gender ratio of each body image was 1:1.

For the body stimuli validation, an independent sample of 35 participants (demographics matched to the main study: age 18–50, 60% female) (Brysbaert, 2019) was recruited via online advertisements distributed through social media platforms. Exclusion criteria mirrored the main study (no IBS, psychiatric/neurological conditions). They rated all body images (bloating, pain, and neutral) using 9-point Likert scales assessing three dimensions: valence (1 = very negative to 9 = very positive), arousal (1 = calm to 9 = highly arousing), and somatic relevance (1 = not at all related to bloating/pain to 9 = extremely related), with Likert-scale ratings (valence, arousal, somatic relevance) aligned with symptom-severity measures (Spiegel et al., 2009). The images were presented in randomized manner to prevent order effects (Lembo et al., 1999; Šolcová & Lačev, 2017). Following data collection, mean ratings were computed separately for each stimulus category (bloating, pain, neutral) to verify that the symptom-related images were indeed perceived as more negative, arousing, and symptom-relevant compared to neutral body images, while also ensuring comparable ratings across neutral stimuli. Mean ratings confirmed significant differences between symptom-related and neutral stimuli (see Results).

Each task comprised 578 trials total, divided equally between 500 ms and 50 ms exposure conditions (289 trials each). Within each exposure duration condition, there were 192 emotional trials (96 congruent/incongruent pairs, balanced for left/right dot location) and 96 neutral trials, plus one initial neutral warm-up trial added at the beginning. In both tasks each trial began with a fixation cross (500 ms), followed by pairs of images presented for 500 ms or 50 ms, with one image appearing to the left and the other to the right of the fixation cross. The 50 ms and 500 ms conditions were completely balanced in terms of trial types and distribution. Notably, while we collected data for both exposure durations (totaling 578 trials per participant), trials with 50 ms stimuli are reserved for separate publication. The current paper focuses exclusively on the 289 trials of 500 ms exposure to examine deliberate attentional allocation.

Threat-related images (in Face Task disgust or anger, in Body Task bloating or abdominal pain) appeared with equal probability on either side. After image offset, a 300 ms blank interval preceded the probe onset. A target dot then appeared for 1000 ms, replacing one of the two images. Participants indicated the dot’s location via button press (left/right). The inter-trial interval (ITI) consisted of a 1000 ms fixation cross. Half of the trials were congruent, and the other half were incongruent. Participants were instructed to indicate the probe’s location by pressing the corresponding button on a response box as quickly and accurately as possible. A fixation cross was displayed for 1000 ms after the probe.

Stimuli were displayed on a 14-inch monitor (1366 × 768 resolution, 60 Hz) using Psychtoolbox in MATLAB. To ensure consistency in low-level visual properties, luminance, contrast, and spatial frequency were matched across stimulus categories using the SHINE toolbox in MATLAB (Willenbockel et al., 2010). This step controlled for potential confounds in attentional allocation driven by basic image features rather than emotional or somatic content. Participants were seated 60 cm from the screen, which was positioned at a 120-degree angle relative to the desktop. All images measured 260 × 300 pixels and were spaced 8 cm apart on a gray background.

### 2.3. Study Design

The study utilized a 2 × 3 × 3 mixed factorial design, combining between-subjects and within-subjects factors. The between-subjects factor was subject type (IBS, individuals with high anxiety and HC), while the within-subjects factors included face type (angry, disgusted, neutral or abdominal pain, bloating and neutral) and dot positioning type (congruent, incongruent, neutral). Reaction time (RT) served as the primary dependent variable.

In congruent trials, the target stimulus (the dot) appeared in the same spatial location as the emotional face (angry/disgusted) or body state (abdominal pain/bloating) within emotional-neutral pairs. Conversely, in incongruent trials, the target stimulus appeared in the opposite location of the emotional stimuli. For neutral trials, pairs consisted of two neutral faces or bodies, with the target stimulus always being neutral.

### 2.4. Procedure

All participants provided signed informed consent and basic demographic information sheet before enrolment in this study, which was approved by the University of Tehran’s ethics committee IR.UT.PSYEDU.1403.018 and conducted in accordance with the Declaration of Helsinki.

After participants completed the STAI and IBS-SSS questionnaires to assess anxiety and IBS symptoms. Upon arrival, they provided informed consent and were guided through a structured protocol. First, they completed pre-task measures, including demographic questionnaires (age, sex, education) and clinical assessments. Participants completed the study in a single laboratory session conducted in a controlled environment with standardized lighting and minimal distractions. Following this, participants received standardized verbal and written instructions for the dot probe tasks, emphasizing speed and accuracy in identifying probe locations. A practice block (10 trials per task) familiarized them with the procedure. After completing the Face Task, participants rested for 2 minutes before proceeding to the Body Task. The Body Task followed an identical trial structure.

The locations of the pictures and the probe were randomized for each participant. The probe disappeared and the next trial was presented if the participant had not responded within 1000 ms. Post-task, participants underwent debriefing to report subjective experiences. Reaction times (RTs) and accuracy were recorded using Psychtoolbox in MATLAB. The time spent on each task was approximately 14 minutes (including instructions).

### 2.5. Data Analysis

Behavioral data from both the face and body tasks were analyzed to quantify attentional biases toward threat-related stimuli (Figure 2 &3). The primary measure was the Bias Index, calculated for each participant as the mean reaction time (RT) on incongruent trials minus the mean RT on congruent trials (Bias Index = RT_incongruent - RT_congruent). Positive values of this index indicate attentional vigilance toward threat (faster responses when the probe replaced threat-related images), while negative values suggest avoidance (slower responses to threat-congruent probes). Trials with RTs < 100 ms (anticipatory responses) or > 3 standard deviations from the participant’s mean RT were excluded as outliers, along with incorrect responses (wrong button pressed). Approximately 90% of trials were retained for analysis after these exclusions.

**Figure 2.**
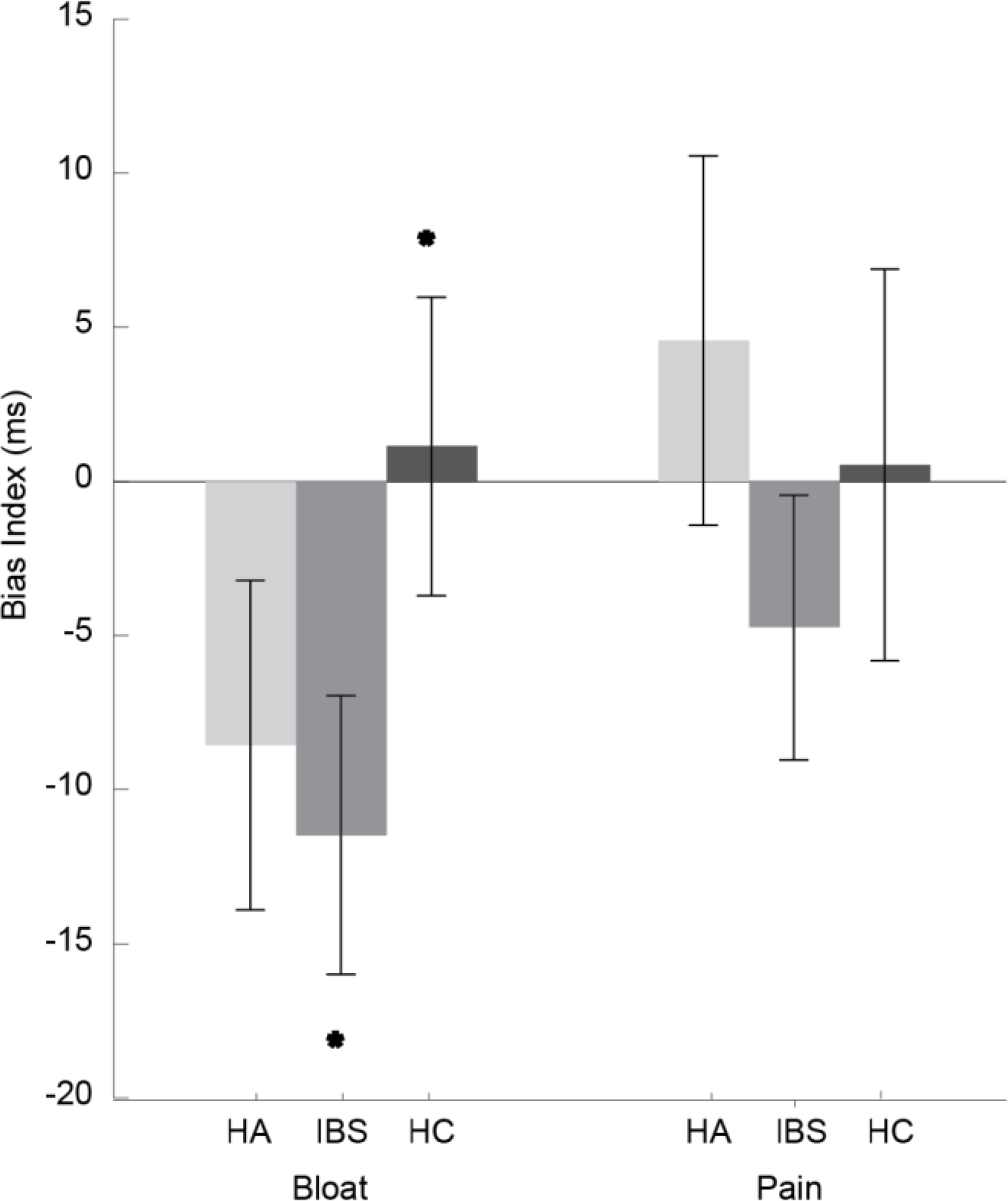
The Bias Index for 500 ms stimuli in the Body Task (Bloating and Pain) across groups with High Anxiety (HA), Irritable Bowel Syndrome (IBS), and Healthy Controls (HC) demonstrated marked differences in attentional patterns. For bloating stimuli, IBS and HA displayed significant negative biases, indicative of avoidance behavior, whereas HC exhibited a subtle positive bias, suggesting engagement with the stimuli. For pain stimuli, HA showed a mild positive bias, IBS demonstrated avoidance with a negative bias, and HC maintained a neutral bias. Error bars represent the standard error of the mean (SEM).

**Figure 3.**
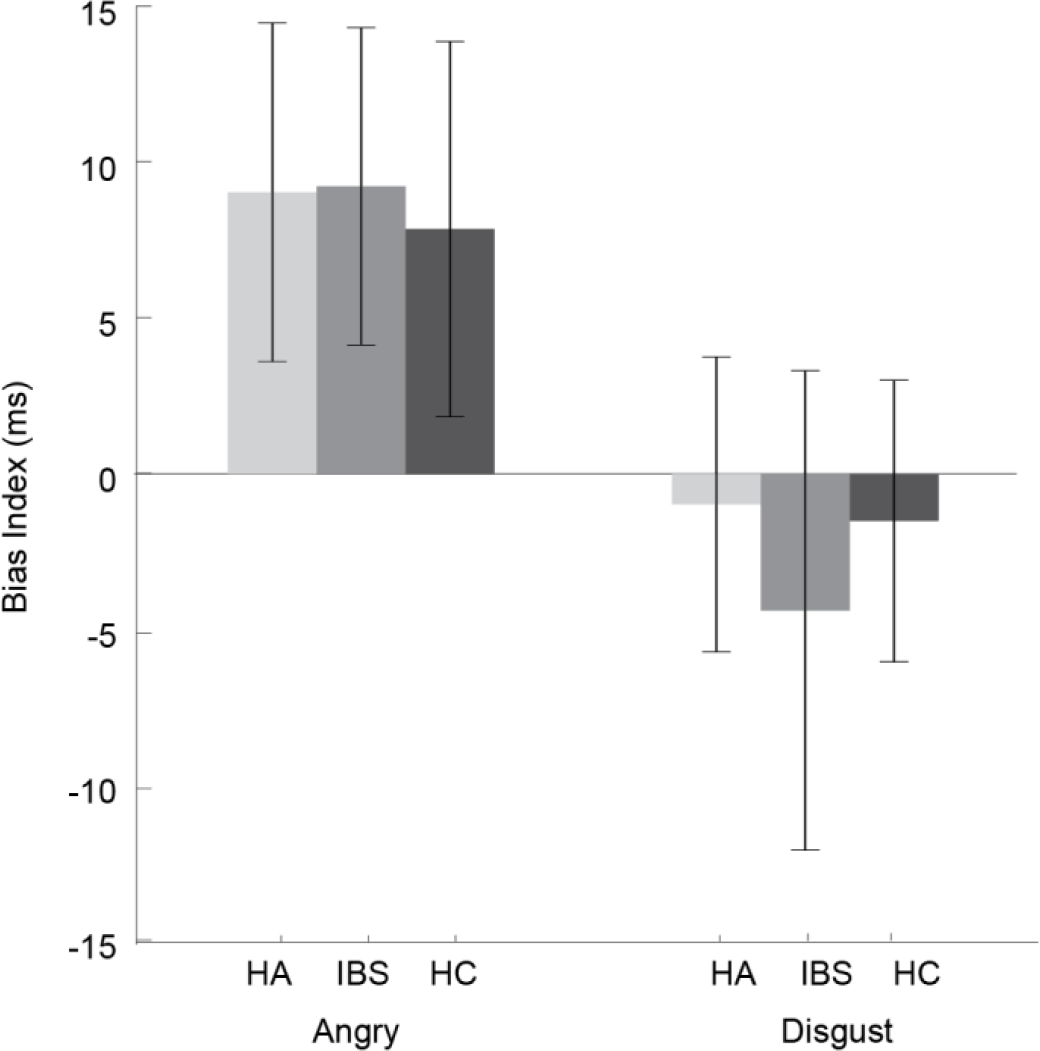
The Bias Index for 500 ms stimuli in Face Task (Angry and Disgust) across High Anxiety (HA), IBS, and Healthy Controls (HC) revealed distinct attentional patterns. All groups demonstrated positive bias scores toward angry faces, indicating a general attentional vigilance to anger-related threat cues, with no statistically significant differences among the groups. In contrast, disgusted faces elicited negative bias indices across all groups, suggesting attentional avoidance without significance difference. Error bars represent standard errors of the mean (SEM).

## 3. Results

Analyses focused on group differences in attentional bias, dot-probe congruency effects, and the moderating roles of anxiety and IBS symptom severity (IBS-SSS). Mixed-design repeated measures ANOVAs, controlling for anxiety and IBS-SSS, were employed with “group” (IBS, HC and HA) as the between-subject factor, “Stimuli” (angry, disgusted and neutral or abdominal pain, bloating and neutral) and “dot type” (congruent versus incongruent) as the within-subject factors. Outcomes also included reaction time (RT) differences, attentional bias indices, and correlations between clinical measures.

### 3.1. Participant Characteristics

The gender, age, and education levels of the group with IBS, group with high anxiety, and healthy control (HC) group were similar (Table 1); however, as expected, significant differences emerged in anxiety scores (p <.001) and IBS symptom severity (IBS-SSS, p <.001). Post-hoc LSD tests revealed that the high anxiety group exhibited substantially higher anxiety levels than HC (mean difference = 32.39, p <.001, Table 1), while no difference was observed between the group with high anxiety and IBS group (p =.970). For IBS-SSS, the IBS group scored markedly higher than both HC (mean difference = 347.69, p <.001) and the group with high anxiety (mean difference = 355.23, p <.001), confirming the clinical distinction of the IBS cohort.

**Table 1.**
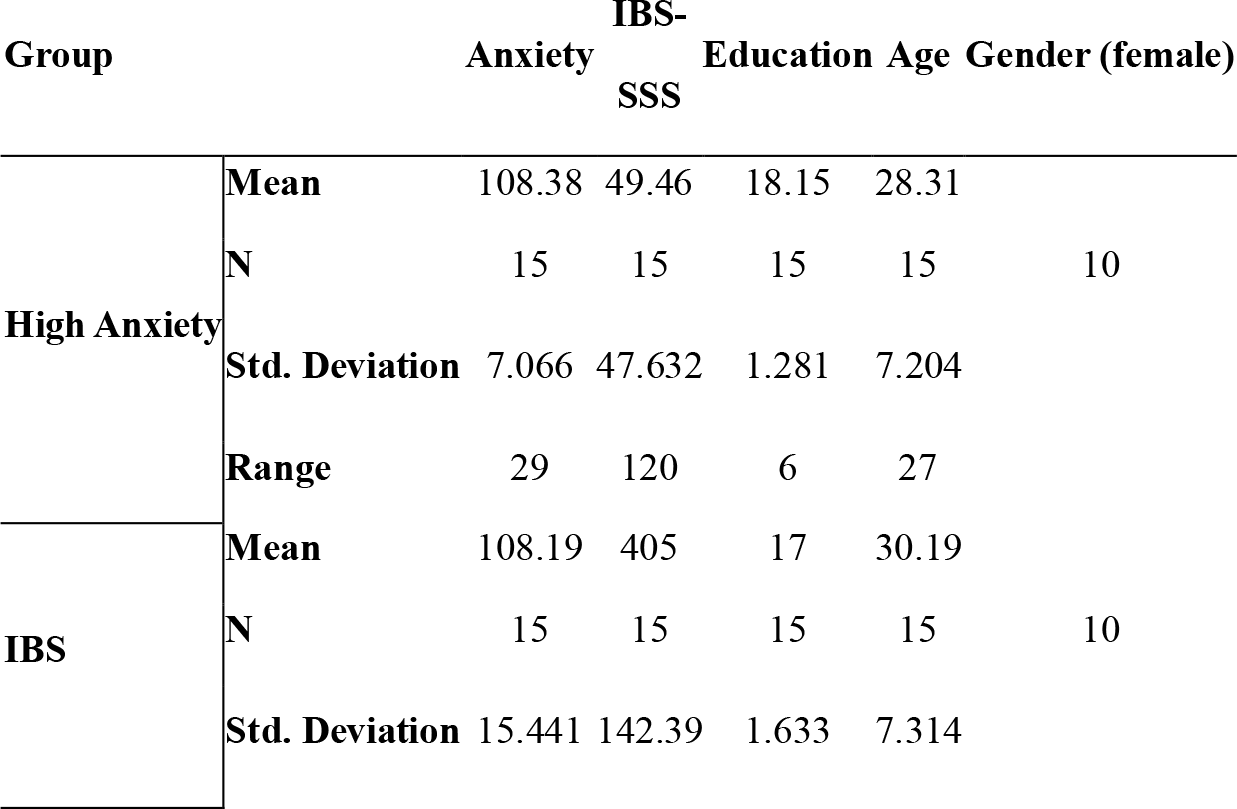
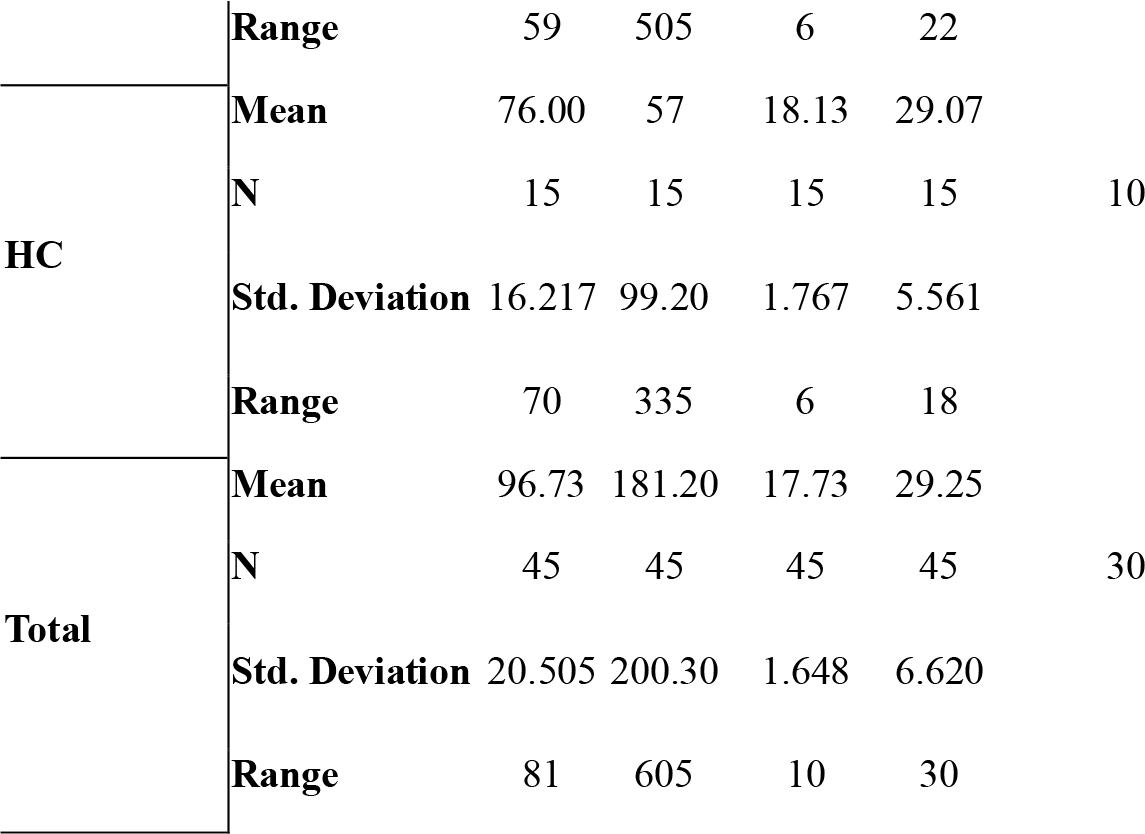
Means and Standard Deviations of Sample Characteristics.

### 3.2. Stimulus Validation Results

Validation data from 35 independent raters confirmed that abdominal pain and bloating images were rated as more negative, arousing, and somatic-relevant than neutral images (all p <.001; see Table 2). Pain and bloating stimuli did not differ significantly from each other (p =.210), supporting their equivalence in threat salience.

**Table 2.** Means and Standard Deviations of Ratings of Body Stimuli (N=35 Independent Raters)

| Stimulus Type | Valence (1–9) | Arousal (1–9) | Somatic Relevance (1–9) | Pairwise Comparisons (p-values) |
| --- | --- | --- | --- | --- |
| Abdominal Pain | 2.1 ± 0.8 | 6.9 ± 1.1 | 8.2 ± 0.9 | Pain vs. Neutral: $p < .001$ |
| Bloating | 2.3 ± 0.7 | 6.7 ± 1.3 | 7.8 ± 1.0 | Bloating vs. Neutral: $p < .001$ |
| Neutral | 5.4 ± 1.0 | 3.1 ± 1.2 | 1.9 ± 0.8 | Pain vs. Bloating: $p = .210$ |

Notes: Valence: 1 = very negative, 9 = very positive; Arousal: 1 = calm, 9 = highly arousing; Somatic Relevance: 1 = not at all related, 9 = extremely related p-values from one-way repeated-measures ANOVAs with Bonferroni correction.

### 3.3. Body Task: Attention Bias of Visceral Stimuli

On Body Task (Figure2), the repeated measures ANOVA revealed significant main effect of Dot Type in the IBS group (*F*(1,13) = 7.55, *p* =.017, η^2^*p* = 0.367 indicating that the IBS group exhibited a structured response pattern to Dot Type variations. A marginal Dot Type × Anxiety interaction was also present in this group (*F*(1,13) = 4.18, *p* =.062, η^2^*p* = 0.243), a significant Group × Stimuli Type interaction (F(2, 2044) = 4.02, p =.018, η^2^ =.004) was also observed, indicating that the effect of stimulus type varied by diagnostic group. Follow-up analyses stratified by group showed that this interaction was primarily driven by the IBS group, which exhibited a significant linear effect of Stimuli Type (F(1, 741) = 7.90, p =.005, η^2^ =.011). In contrast, neither the HA (F(1, 605) = 0.65, p =.422, η^2^ =.001) nor the HC (F(1, 698) = 2.13, p =.145, η^2^ =.003) groups showed significant linear trends.

Consistent with the within-subjects’ findings, pairwise comparisons revealed that patients with IBS differed significantly from HC in response to bloat stimuli (Mean Difference = -12.83 ms, SE = 5.01, p =.013, d = 0.61), with patients with IBS showing heightened reactivity. A marginal trend suggested that individuals with HA responded less strongly than HC to bloat stimuli (Mean Difference *= -8.92 ms, SE = 5.24, p =*.*094, d = 0.42*), though this did not reach significance. No group differences were observed for pain stimuli (all p >.190). A marginal Dot Type × Anxiety interaction was also present in this group (*F*(1,13) = 4.18, *p* =.062, η^2^*p* = 0.243), suggesting that anxiety levels may influence dot processing in IBS.

No significant effects were found for Dot Type (all p >.900) or the Dot Type × Stimuli Type interaction (all p >.380), indicating that the dot pattern manipulation did not influence task performance. In the group with HA, no significant main effects or interactions involving Dot Type or Stimuli Type were observed (all *p* >.342). After controlling anxiety and IBS symptom severity, the Dot Type effect remained significant in the IBS group (*p* =.017). No significant relationships were found between anxiety or IBS-SSS and Stimuli Type processing in any group (all *p* >.419). A between-subjects ANOVA was performed for neutral trials; the result revealed no significant effects between three groups for neutral trials (F(2,39) = 0.84, p =.441).

### 3.4. Face Task: Attention Bias of Emotional Faces

In Face Task (Figure3), the analysis revealed a marginally significant main effect of Dot Type (*F*(1,39) = 3.52, *p* =.068, η^2^p = 0.083), suggesting a trend toward differential attentional allocation between congruent and incongruent trials. Further within-group analyses demonstrated significant Dot Type effects in both the IBS (*F*(1,13) = 7.45, *p* =.017, η^2^p = 0.364) and HC groups (*F*(1,12) = 6.03, *p* =.030, η^2^p = 0.334), indicating attentional bias in these groups. Notably, anxiety levels significantly modulated Dot Type effects in IBS (*F*(1,13) = 7.79, *p* =.015, η^2^p = 0.375) and HC participants (*F*(1,12) = 8.38, *p* =.013, η^2^p = 0.411), highlighting anxiety’s influence on attentional processing.

No significant main effect of Stimuli Type (*F*(1,39) = 2.57, *p* =.117, η^2^p = 0.062) or interactions involving Stimuli Type and Anxiety (*F*(1,39) = 3.18, *p* =.082, η^2^p = 0.075) or IBS-SSS (*F*(1,39) = 2.22, *p* =.144, η^2^p = 0.054) were observed. Similarly, the three-way interaction between Dot Type, Stimuli Type, and Group was non-significant (*F*(2,39) = 1.51, *p* =.234, η^2^p = 0.072). When controlling for Anxiety and IBS-SSS, the overall effect of Group on Dot Type modulation remained non-significant (*F*(2,39) = 1.43, *p* =.251, η^2^p = 0.068). Between-subjects ANOVA for neutral trials revealed no significant main effects of group (High Anxiety, IBS, HC) on reaction times (all p >.05). Pairwise comparisons showed marginally faster responses in healthy controls (HC) compared to individuals with HA and patients with IBS (mean difference = 44.87 ms, p =.07).

### 3.5. Attention Bias Index Analysis

Subsequently, the attentional bias index was calculated by subtracting the mean reaction time (RT) of valid cues from the mean RT of invalid ones. To investigate the specific group differences in attentional biases, post-hoc comparisons were conducted on the bias indices (Table 3). For bloating-related stimuli, patients with IBS showed significant avoidance compared to healthy controls (mean bias index difference = -12.83 ms, SE = 5.01, *p* =.013, *d* = 0.61), while the difference between HA and HC participants approached marginal significance (mean difference = -8.92 ms, SE = 5.24, *p* =.094, *d* = 0.42). The comparison between HA and IBS groups was non-significant (mean difference = 3.85 ms, SE = 5.12, *p* =.456, *d* = 0.18).

**Table 3.** Post-Hoc Comparisons of Attentional Bias Indices Between Groups.

| Stimulus Type | Comparison | Mean Difference (ms) | SE | P-value | 95% CI | Cohen's d |
| --- | --- | --- | --- | --- | --- | --- |
| **Bloat Stimuli** | High Anxiety vs IBS | 3.85 | 5.12 | 0.456 | [-6.52, 14.22] | 0.18 |
|  | High Anxiety vs HC | -8.92 | 5.24 | 0.094 | [-19.50, 1.66] | 0.42 |
|  | IBS vs HC | -12.83 | 5.01 | 0.013 | [-22.92, -2.74] | 0.61 |
| **Pain Stimuli** | High Anxiety vs IBS | 8.15 | 6.22 | 0.194 | [-4.42, 20.72] | 0.32 |
|  | High Anxiety vs HC | 4.92 | 6.35 | 0.441 | [-7.92, 17.76] | 0.19 |
|  | IBS vs HC | -3.23 | 4.87 | 0.51 | [-13.07, 6.61] | 0.15 |
| **Angry Faces** | High Anxiety vs IBS | -0.19 | 7.85 | 0.981 | [-16.04, 15.66] | 0.01 |
|  | High Anxiety vs HC | 1.16 | 7.97 | 0.885 | [-14.92, 17.25] | 0.06 |
|  | IBS vs HC | 1.36 | 7.55 | 0.858 | [-13.90, 16.61] | 0.07 |
| **Disgusted Faces** | High Anxiety vs IBS | 3.38 | 8.57 | 0.696 | [-13.93, 20.69] | 0.13 |
|  | High Anxiety vs HC | 7.45 | 8.7 | 0.397 | [-10.12, 25.02] | 0.43 |
|  | IBS vs HC | -2.85* | 7.25 | 0.695 | [-17.35, 11.65] | -0.17 |

For pain-related stimuli, individuals with HA exhibited a non-significant tendency toward vigilance relative to patients with IBS (mean difference = 8.15 ms, SE = 6.22, *p* =.194, *d* = 0.32) and HC (mean difference = 4.92 ms, SE = 6.35, *p* =.441, *d* = 0.19). Patients with IBS demonstrated slight, non-significant avoidance compared to controls (mean difference = -3.23 ms, SE = 4.87, *p* =.510, *d* = 0.15).

Processing of facial stimuli showed no significant group differences. For angry faces, all comparisons were non-significant (all *p* >.85, *d* = 0.01-0.07). Disgusted faces elicited a medium effect-sized difference between participants with HA and HC (mean difference = 7.45 ms, SE = 8.70, *p* =.397, *d* = 0.43), though this did not reach statistical significance. Patients with IBS displayed mild avoidance of disgusted faces relative to controls (mean difference = -2.85 ms, SE = 7.25, *p* =.695, *d* = -0.17), while the HA vs IBS comparison was non-significant (mean difference = 3.38 ms, SE = 8.57, *p* =.696, *d* = 0.13).

### 3.6. Correlations Between IBS Symptom Severity and Anxiety

A significant positive correlation was found between IBS symptom severity (IBS-SSS) and anxiety (STAI scores) in the IBS group (r = 0.574, p =.020, N = 15), indicating that participants with more severe IBS symptoms reported higher levels of anxiety. This association was not observed in either the HA group (r = 0.057, p =.853, N = 15) or HC (r = 0.386, p =.156, N = 15). No significant correlations were found between anxiety scores and overall reaction times in any group: HA (r = -0.03, p =.93), IBS (r = -0.31, p =.24), or HC (r = -0.34, p =.21).

### 3.7. Group Differences in Overall Reaction Times

The analysis of mean reaction times (RTs) across the three groups—HC, High Anxiety, and IBS— revealed significant differences. As illustrated in figure 4, the HC group exhibited the lowest mean RT (399.57 ± 12.61 ms), followed by the HA group (421.84 ± 14.32 ms), and the IBS group, which displayed the highest mean RT (430.28 ± 10.87 ms). Post-hoc comparisons using Tukey’s HSD test revealed significant differences in reaction times between groups. HC group demonstrated significantly faster RTs compared to individuals with IBS group across most task conditions (p =.024) with the exception of incongruent neutrals trials of Body Task where no significant difference emerged. While the HA group exhibited numerically slower RTs relative to HC, this difference was not statistically significant (p =.375). Similarly, although patients with IBS showed marginally slower RTs than the group with HA, this contrast did not reach statistical significance (p =.133).

**Figure 4.**
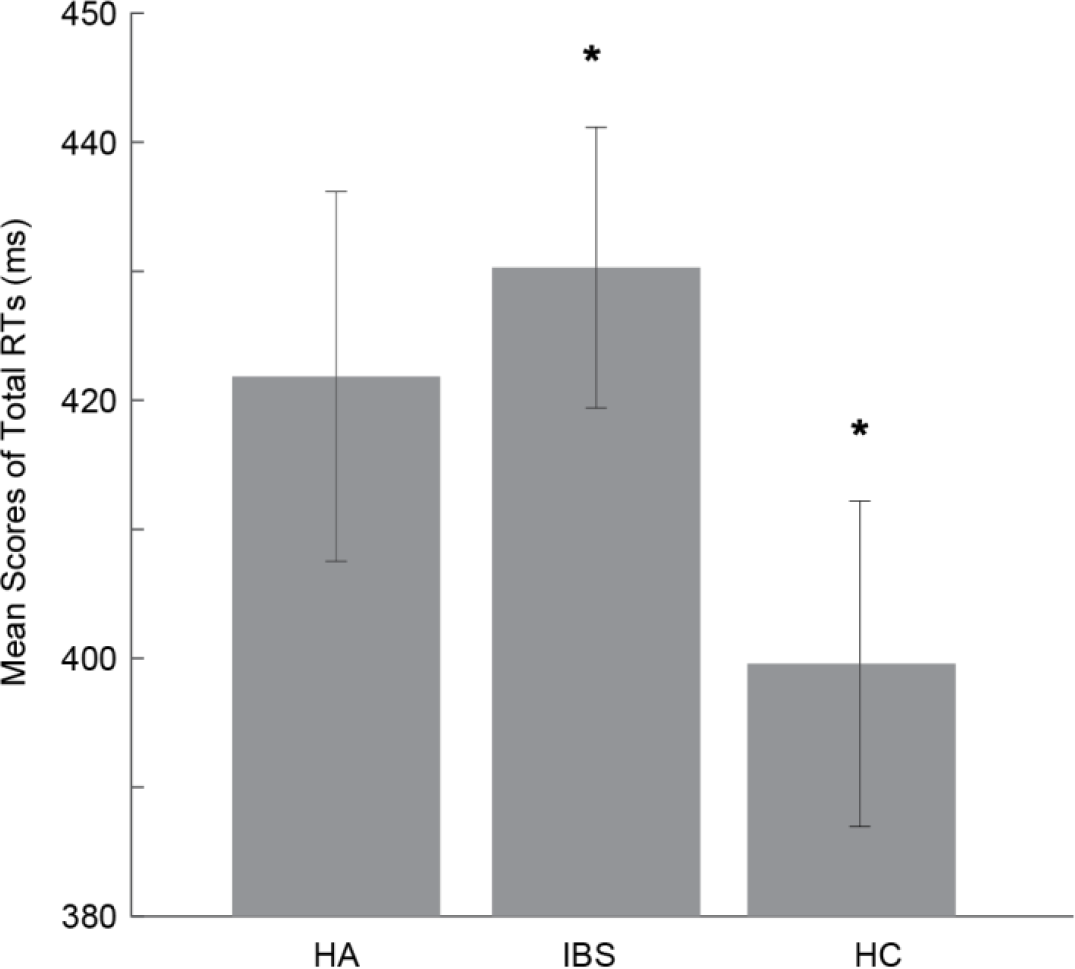
Mean total reaction times (RTs) across study groups (High Anxiety, IBS, and Healthy Controls). Error bars represent standard deviations. The asterisk (*) indicates a statistically significant difference between IBS and Healthy Controls (*p* <.05). High Anxiety participants showed intermediate RTs that did not significantly differ from either group.

## 4. Discussion

The current study investigated attentional bias toward somatic-relevant (bloating and pain) and emotional (angry and disgusted faces) stimuli in individuals with irritable bowel syndrome (IBS), individuals with high anxiety (HA), and healthy controls (HC). Using a dot-probe paradigm across body- and face-related tasks, we examined the influence of anxiety and symptom severity on attentional patterns, while controlling group-level differences.

In the Body Task, patients with IBS exhibited a significant main effect of dot allocation, demonstrating heightened responses to incongruent versus congruent stimuli. The marginal interaction congruent/incongruent and anxiety further implies that anxiety levels may modulate this attentional bias, a notion supported by the significant correlation between IBS symptom severity and anxiety within the IBS group. Notably, the HA group did not exhibit significant attentional biases in the Body Task, indicating that the observed effects are more closely associated with IBS symptomatology than with anxiety alone.

Interestingly, avoidance of pain-related stimuli was less pronounced and did not reach statistical significance across groups. This finding contrasts with some studies suggesting that chronic pain patients display attentional biases toward pain-related information (Schoth et al., 2019; Todd et al., 2018). The absence of significant avoidance in our study may be attributed to the specific nature of IBS, where bloating is a more prevalent and bothersome symptom than pain (Kanazawa et al., 2016; Schmulson & Drossman, 2017). Consequently, attentional biases in IBS may be more closely associated with bloating-related cues than pain-related ones.

The Face Task revealed a marginally significant main effect of dot allocation across all participants, with significant within-group effects in both the IBS and HC groups. This indicates that both groups allocated attention differentially between congruent and incongruent trials. The significant modulation of Dot Type effects by anxiety levels in these groups underscores the influence of anxiety on attentional processing of facial stimuli. However, the HA group did not demonstrate significant main effects or interactions, suggesting that high anxiety alone does not account for the observed attentional biases.

Contrary to expectations, the HA group did not exhibit significant main effects or interactions in either task. This absence of detectable attentional biases may stem from several factors. First, the heterogeneity of anxiety-related attentional biases is well-documented (Bantin et al., 2016); individuals with high anxiety may display vigilance toward threat, avoidance, or no bias at all (Ioannou et al., 2004; Koster et al., 2006). Such variability can obscure group-level effects. Second, methodological considerations, such as the nature of the stimuli or task parameters, might influence the detection of biases. For instance, some studies suggest that attentional biases in anxious individuals are more pronounced under specific conditions or with particular stimuli (MacLeod et al., 2019). These factors collectively suggest that the dot-probe task may not consistently capture attentional biases in high-anxiety individuals.

The findings revealed a significant avoidance of bloating-related stimuli in the IBS group relative to HC, as evidenced by a negative bias index (mean difference = -12.83 ms, *p* =.013, *d* = 0.61). This negative index indicates slower responses to threat-congruent probes (i.e., delayed engagement with bloat images), aligning with cognitive-behavioral models of somatic symptom avoidance in IBS (Hesser et al., 2018). This pattern of attentional disengagement suggests that IBS patients actively avoid visceral threat cues, potentially to preempt anticipated discomfort—a behavior consistent with negative self-schema and low subjective vigor, as highlighted by Phillips et al. (2014). Nadinda et al. (2024) attribute IBS avoidance to imbalanced predictive coding: hyper-precise negative expectancies (e.g., anticipating bloating) outweigh low-precision sensory input, creating a self-reinforcing cycle that sustains symptoms. Notably, this contrasts with the initial interpretation of vigilance (Kilpatrick et al., 2010; Posserud et al., 2009), underscoring that IBS patients preferentially disengage from gastrointestinal threat cues rather than fixating on them.

Notably, the avoidance was specific to bloating stimuli. Despite being another key symptom domain in IBS, pain-related stimuli did not elicit significant avoidance in any group. Instead, there was a non-significant trend toward vigilance in the HA group, and mild, non-significant avoidance in IBS. This finding may reflect the more generalized threat value of pain, which could provoke increased attention even in non-somatic anxiety, as opposed to bloating which may be more uniquely associated with IBS-specific distress. Furthermore, the weaker behavioral differentiation across groups in response to pain cues—combined with larger error bars (as shown in Figure 2)— suggests greater interindividual variability in how pain-related threat is processed.

With respect to emotional facial expressions, no significant group differences were found, though subtle patterns emerged. All groups showed a mild, nonsignificant bias toward vigilance for angry faces. This pattern is consistent with a general attentional preference for threat-related expressions, which may be evolutionarily adaptive and not specific to clinical anxiety or IBS.

In the Face Task, the bias indices for disgusted faces were negative across all groups, suggesting a general tendency for attentional avoidance. The IBS group showed the strongest avoidance, while the High Anxiety (HA) and Healthy Control (HC) groups showed less avoidance. Despite this pattern, no significant differences were found between the groups. Though these effects did not reach statistical significance (all *p* >.39), the directionality may be relevant. Disgust, as a social-emotional cue often linked to contamination or social rejection, may resonate more with individuals experiencing somatic symptoms (e.g., IBS) due to heightened interoceptive sensitivity or self-consciousness. However, the lack of significant effects limits the strength of this interpretation and suggests that facial expressions may not have been potent enough triggers in this context compared to symptom-relevant bodily cues. Research indicates a positive correlation between disgust sensitivity and IBS symptoms. A study involving healthy individuals found that higher disgust sensitivity was associated with more severe IBS symptoms (Formica et al., 2022). This association suggests that individuals with heightened sensitivity to disgust may be more prone to experiencing IBS-related discomfort.

An important pattern observed across tasks is that while the IBS and HC groups showed significant within-group attentional effects, the HA group did not. The absence of significant main effects or interactions in the HA group suggests that their attentional deployment may be more diffuse or less stimulus-specific. In contrast to the IBS group, whose attentional patterns were likely shaped by chronic somatic experience, and the HC group, which may exhibit normative attentional patterns in response to social-emotional cues, the HA group might show greater variability in attentional responses, lacking a clear threat schema tied to either bodily or emotional stimuli. This aligns with the possibility that attentional bias in anxiety is contextually modulated and may require task parameters that better capture hypervigilance to generalized threat cues (MacLeod et al., 2019).

A significant positive correlation was observed between IBS symptom severity and anxiety levels within the IBS group, suggesting that more severe IBS symptoms are associated with higher anxiety. This relationship was not evident in the HA or HC groups, highlighting a unique interplay between symptom severity and anxiety in patients with IBS (Hubbard et al., 2015).

Analysis of reaction times (RTs) revealed that HC participants responded significantly faster than patients with IBS across most task conditions, with the exception of incongruent neutral trials in the Body Task. While the HA group exhibited numerically slower RTs compared to HC, these differences were not statistically significant. These findings suggest that patients with IBS may experience generalized cognitive slowing or increased cognitive load, potentially due to the chronic nature of their symptoms (Aizawa et al., 2012; Phillips et al., 2014). This is supported by studies that have found hypoactivity in the default mode network (DMN) of IBS patients at resting state, which may reflect a state of continuous hypervigilance and spontaneous pain perception (Nisticò et al., 2022).

### 4.1. Implications

The observed attentional biases toward gastrointestinal (GI)-related stimuli in patients with IBS suggest that cognitive-behavioral interventions should focus on modifying these biases. Techniques such as attention bias modification training could be employed to redirect attention away from symptom-related cues, potentially reducing symptom severity and improving quality of life (Tayama et al., 2018).

Furthermore, the correlation between heightened disgust sensitivity and IBS symptom severity indicates the need for therapeutic strategies that specifically address emotional responses to disgust. Interventions could include exposure therapy (Axelsson et al., 2023) or cognitive restructuring to alter maladaptive disgust reactions(Fink et al., 2018), thereby alleviating IBS symptoms. Recognizing the role of personality traits and emotional patterns in IBS underscores the importance of personalized psychological interventions. Tailoring treatments to individual cognitive-affective profiles may enhance therapeutic outcomes.

The interplay between psychological factors and GI symptoms in patients with IBS calls for a multidisciplinary treatment approach (Chiou & Nurko, 2010). Collaboration among gastroenterologists, psychologists, and other healthcare professionals is essential to address both the physiological and psychological aspects of IBS effectively. Integrating cognitive-behavioral strategies aimed at attentional biases and emotional responses, particularly disgust sensitivity, into comprehensive treatment plans may improve patient outcomes in IBS. Future research with larger cohorts is necessary to elucidate the relationship between disgust sensitivity and attentional biases in patients with IBS. Additionally, exploring the neural mechanisms underlying these biases could provide deeper insights into the cognitive-affective profile of individuals with IBS.

### 4.2. Limitations and Future Directions

The current study provides important insights into attentional biases in IBS, but several limitations should be acknowledged. First, while our sample size was sufficient to detect large effects—such as the significant avoidance of bloating-related stimuli in IBS patients—it may have been underpowered to identify more subtle group differences or interactions. Future studies would benefit from larger samples to enhance statistical power and better capture nuanced effects.

Second, although we carefully validated our body stimuli, the images represented a limited range of demographics. Expanding stimulus diversity in future workcould improve the generalizability of findings.

Third, the absence of significant effects in the high-anxiety group raises questions about whether the selected stimuli adequately captured threat relevance for non-IBS anxiety. Future research could compare responses to IBS-specific versus generalized anxiety cues to clarify this distinction.

Additionally, the dot-probe task’s reliability issues suggest that alternative or complementary measures, such as eye-tracking or neuroimaging, could provide more robust assessments of attentional biases.

## 5. Conclusion

This study revealed distinct attentional bias patterns in patients with IBS patients, characterized by significant avoidance of bloating-related stimuli, which was modulated by anxiety levels. In contrast, high anxiety alone did not produce consistent attentional biases, suggesting that IBS-related attentional patterns are uniquely tied to somatic symptom processing. The correlation between IBS symptom severity and anxiety underscores the interplay between cognitive and gastrointestinal dysfunction.

## Ethics Approval and Consent to Participate

This study was approved by the Ethics Committee of the University of Tehran (IR.UT.PSYEDU.1403.018). All procedures adhered to the Declaration of Helsinki. Written informed consent was obtained from all participants prior to enrolment.

## Consent for Publication

Not applicable (no individual personal data/images are published).

## Availability of Data and Materials

The data are not publicly available to protect participant confidentiality. The datasets used and/or analyzed during the current study are, however, available from the corresponding author on reasonable request and with the permission from the University of Tehran.

## Acknowledgements

The authors would like to thank all participants for taking part in the experiment.

## Authors’ Information

Reyhaneh Akbari: PhD Candidate, Health Psychology, University of Tehran.

Fateme Dehghani-Arani: Assistant Professor, Psychology, University of Tehran.

Mohsen Honar: M.Sc. in Cognitive Sciences, University of Tehran.

Nazila Shahmansouri: Associate Professor, Psychosomatic Medicine, Tehran University of Medical Sciences

Ehsan Rezayat: Assistant Professor, Cognitive Neuroscience, University of Tehran.

